## Supplementary Materials for "Testing microbiome associations with censored survival outcomes at both the community and individual taxon levels"

**Table S1.** Type I error of the global tests for simulated data in other cases

| Censoring | $n$ | Scenario | $\beta_{XZ}$ | LDM- | | | permanovaFL- | | | MiRKAT-S | OMiSA |
|---|---|---|---|---|---|---|---|---|---|---|---|
| | | | | c | m | d | c | m | d | | |
| 75% | 100 | M1 | 0 | 0.050 | 0.049 | 0.053 | 0.051 | 0.051 | 0.052 | 0.051 | 0.053 |
| | | | 0.8 | 0.052 | 0.050 | 0.048 | 0.050 | 0.050 | 0.050 | 0.032 | 0.038 |
| | | | 0.8* | 0.332 | 0.329 | 0.314 | 0.239 | 0.228 | 0.228 | 0.233 | 0.261 |
| | | M2 | 0 | 0.048 | 0.047 | 0.049 | 0.049 | 0.048 | 0.050 | 0.051 | 0.052 |
| | | | 0.8 | 0.051 | 0.053 | 0.049 | 0.050 | 0.050 | 0.047 | 0.009 | 0.038 |
| | | | 0.8* | 0.469 | 0.480 | 0.442 | 0.475 | 0.473 | 0.437 | 0.474 | 0.261 |
| 25% | 100 | M1 | 0 | 0.052 | 0.051 | 0.052 | 0.052 | 0.051 | 0.051 | 0.053 | 0.052 |
| | | | 0.8 | 0.047 | 0.044 | 0.048 | 0.049 | 0.046 | 0.049 | 0.030 | 0.033 |
| | | | 0.8* | 0.806 | 0.812 | 0.725 | 0.624 | 0.628 | 0.593 | 0.643 | 0.697 |
| | | M2 | 0 | 0.043 | 0.045 | 0.044 | 0.045 | 0.044 | 0.045 | 0.047 | 0.052 |
| | | | 0.8 | 0.049 | 0.047 | 0.046 | 0.049 | 0.050 | 0.047 | 0.009 | 0.033 |
| | | | 0.8* | 0.912 | 0.919 | 0.86 | 0.925 | 0.931 | 0.864 | 0.931 | 0.694 |
| 50% | 50 | M1 | 0 | 0.045 | 0.046 | 0.044 | 0.045 | 0.046 | 0.044 | 0.051 | 0.047 |
| | | | 0.8 | 0.045 | 0.047 | 0.047 | 0.049 | 0.050 | 0.048 | 0.032 | 0.027 |
| | | | 0.8* | 0.335 | 0.327 | 0.294 | 0.218 | 0.217 | 0.207 | 0.225 | 0.304 |
| | | M2 | 0 | 0.042 | 0.044 | 0.043 | 0.047 | 0.046 | 0.046 | 0.050 | 0.047 |
| | | | 0.8 | 0.044 | 0.045 | 0.040 | 0.046 | 0.047 | 0.044 | 0.006 | 0.027 |
| | | | 0.8* | 0.462 | 0.471 | 0.435 | 0.474 | 0.477 | 0.445 | 0.489 | 0.304 |

Note: See the note to Table 1. All event times here were simulated from the Cox model.

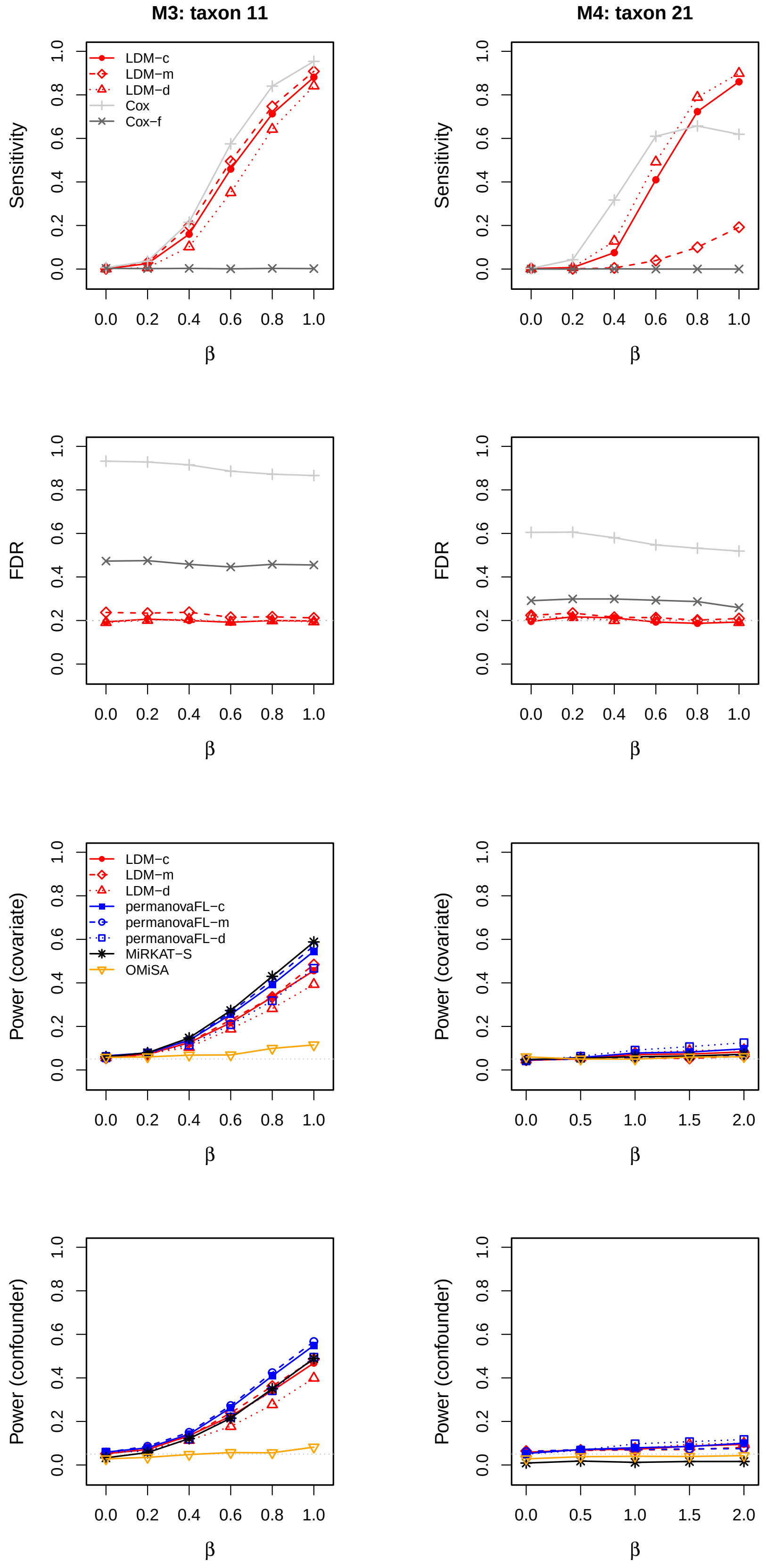


Figure S1: Results in scenarios M3 and M4 when taxa 11 and 21, respectively, were associated with the event time. Results of sensitivity and empirical FDR were obtained when $X_i$ was a confounder ($\beta_{XZ} = 0.8$).

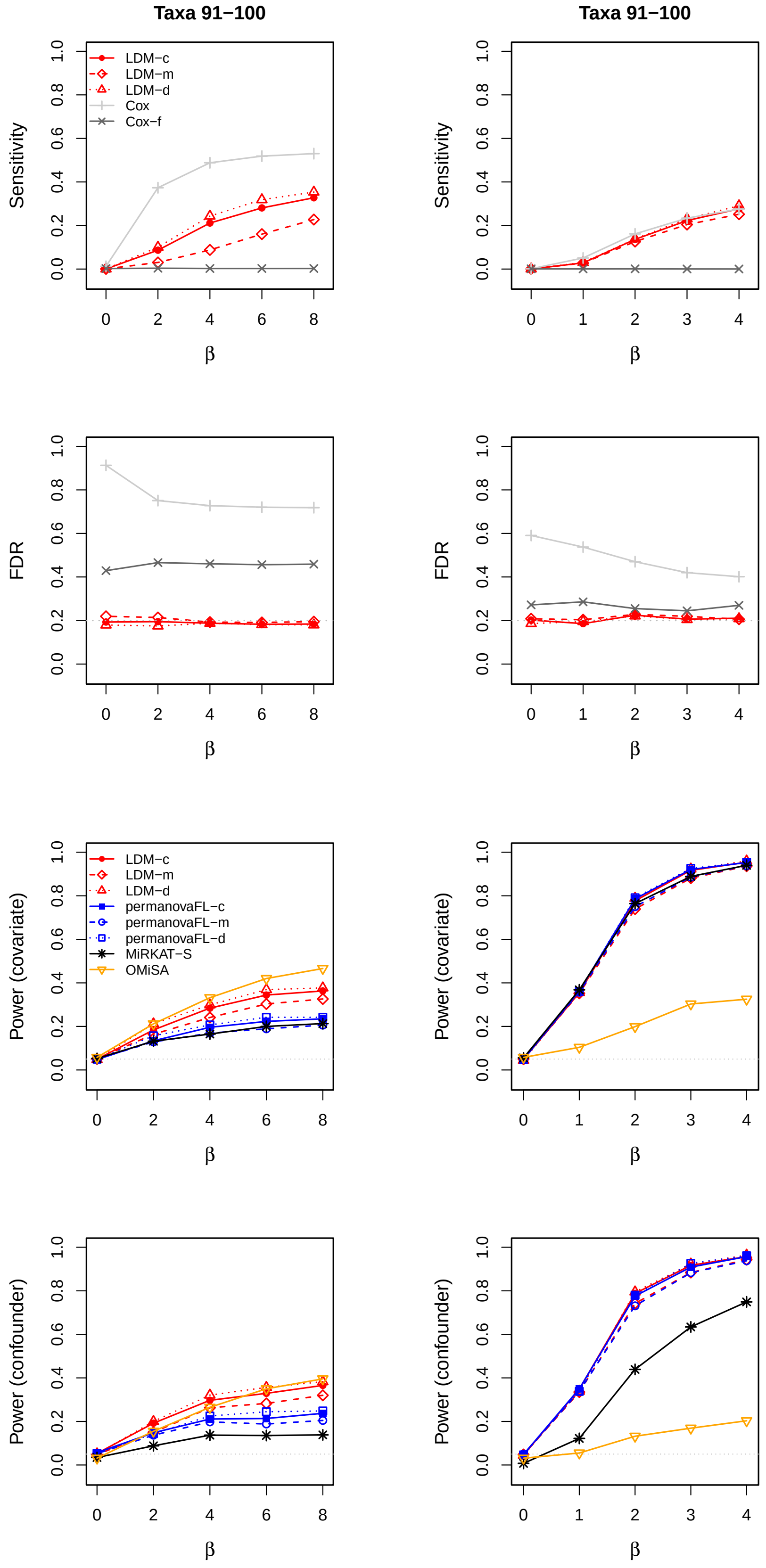


Figure S2: Results in the scenario when rare taxa (taxa 91–100) were associated with the event time. Left column: data were simulated and analyzed based on the relative abundance scale, same as in model M1. Right column: data were simulated and analyzed based on the presence-absence scale (except for OMiSA), same as in model M2. The censoring rate was 50% and $n = 100$. Results of sensitivity and empirical FDR were obtained when $X_i$ was a confounder ($\beta_{XZ} = 0.8$).

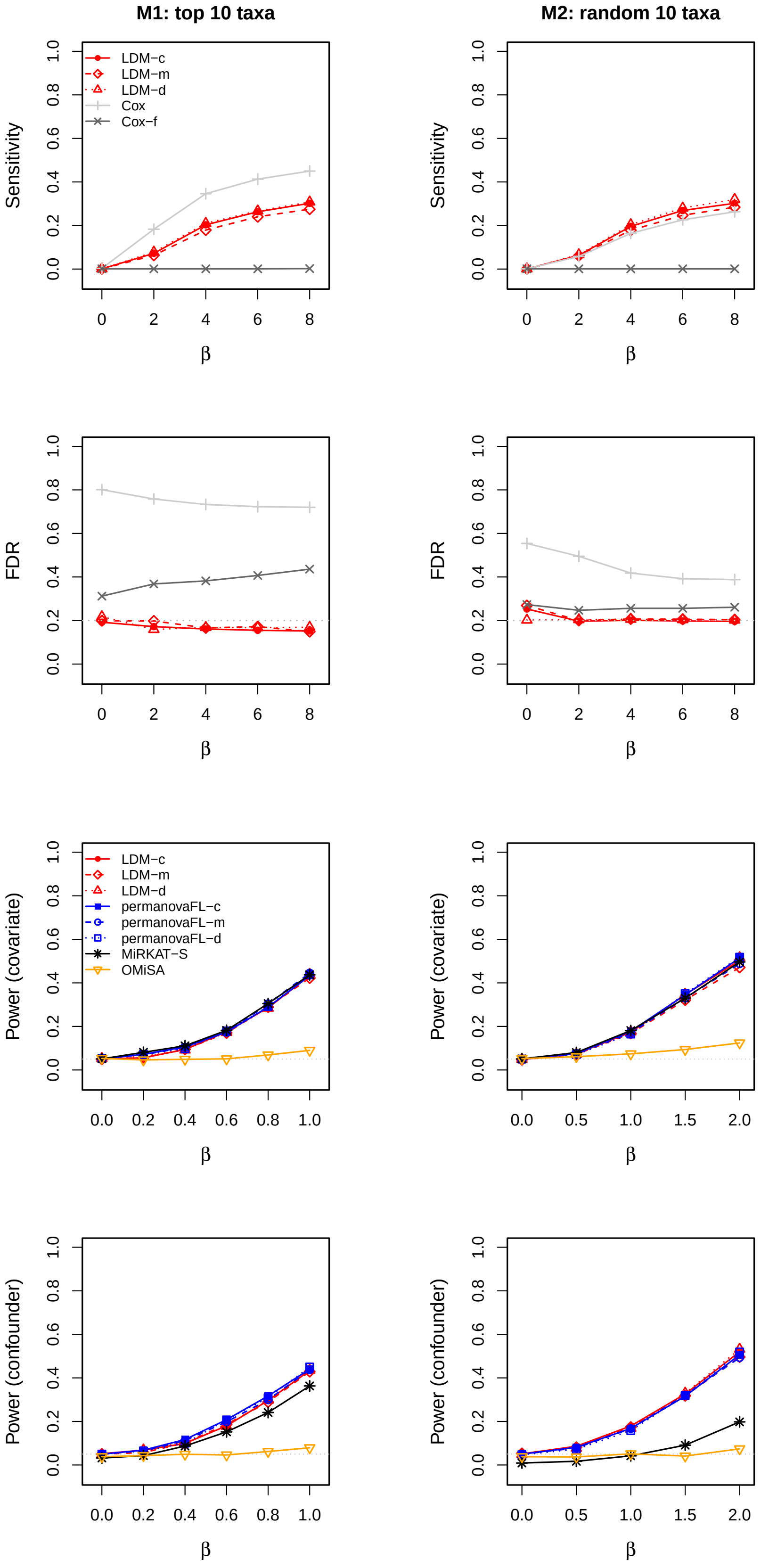


Figure S3: Results for simulated data with 75% censoring and $n = 100$. Results of sensitivity and empirical FDR were obtained when $X_i$ was a confounder ($\beta_{XZ} = 0.8$).

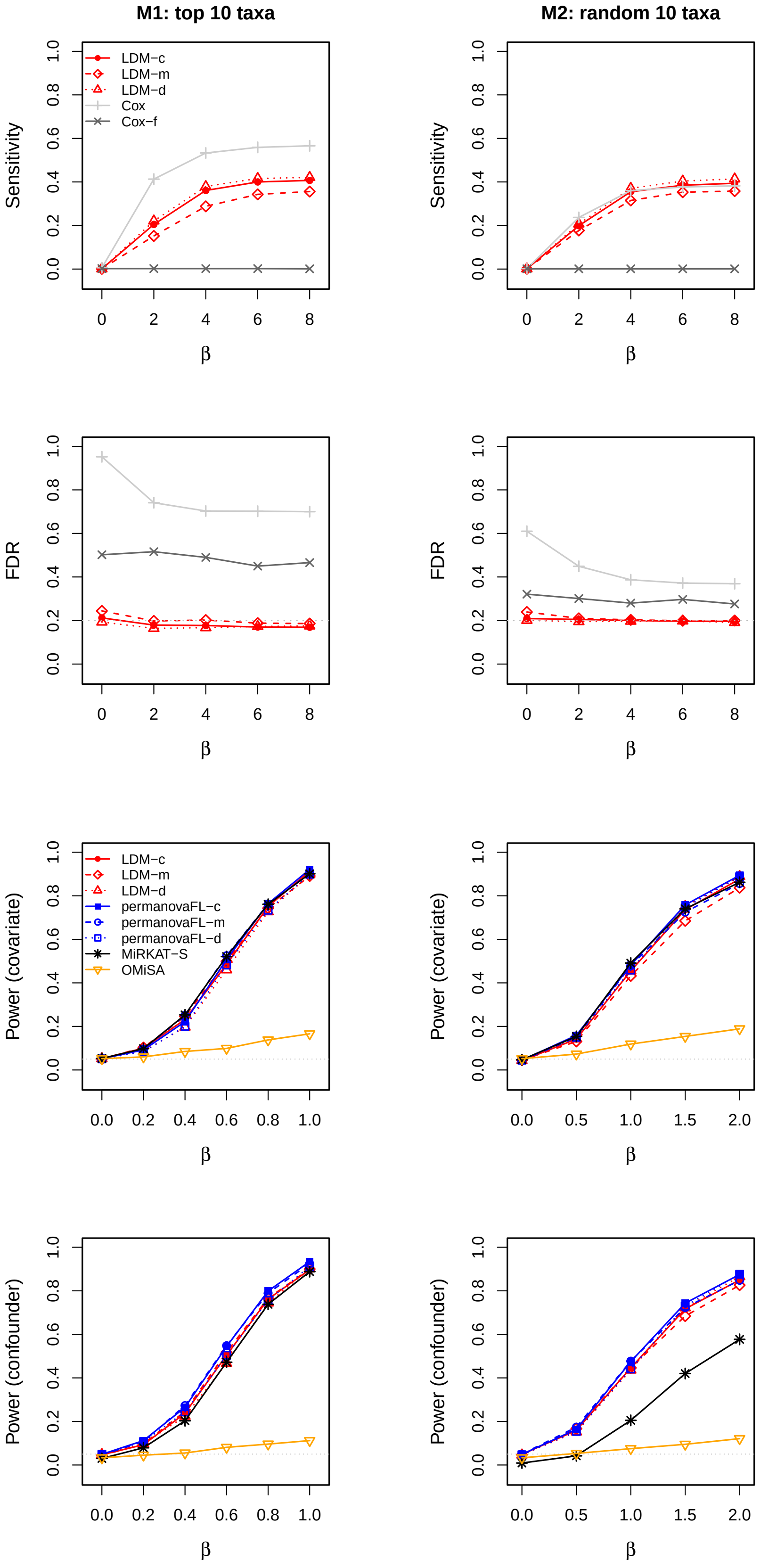


Figure S4: Results for simulated data with 25% censoring and $n = 100$. Results of sensitivity and empirical FDR were obtained when $X_i$ was a confounder ($\beta_{XZ} = 0.8$).

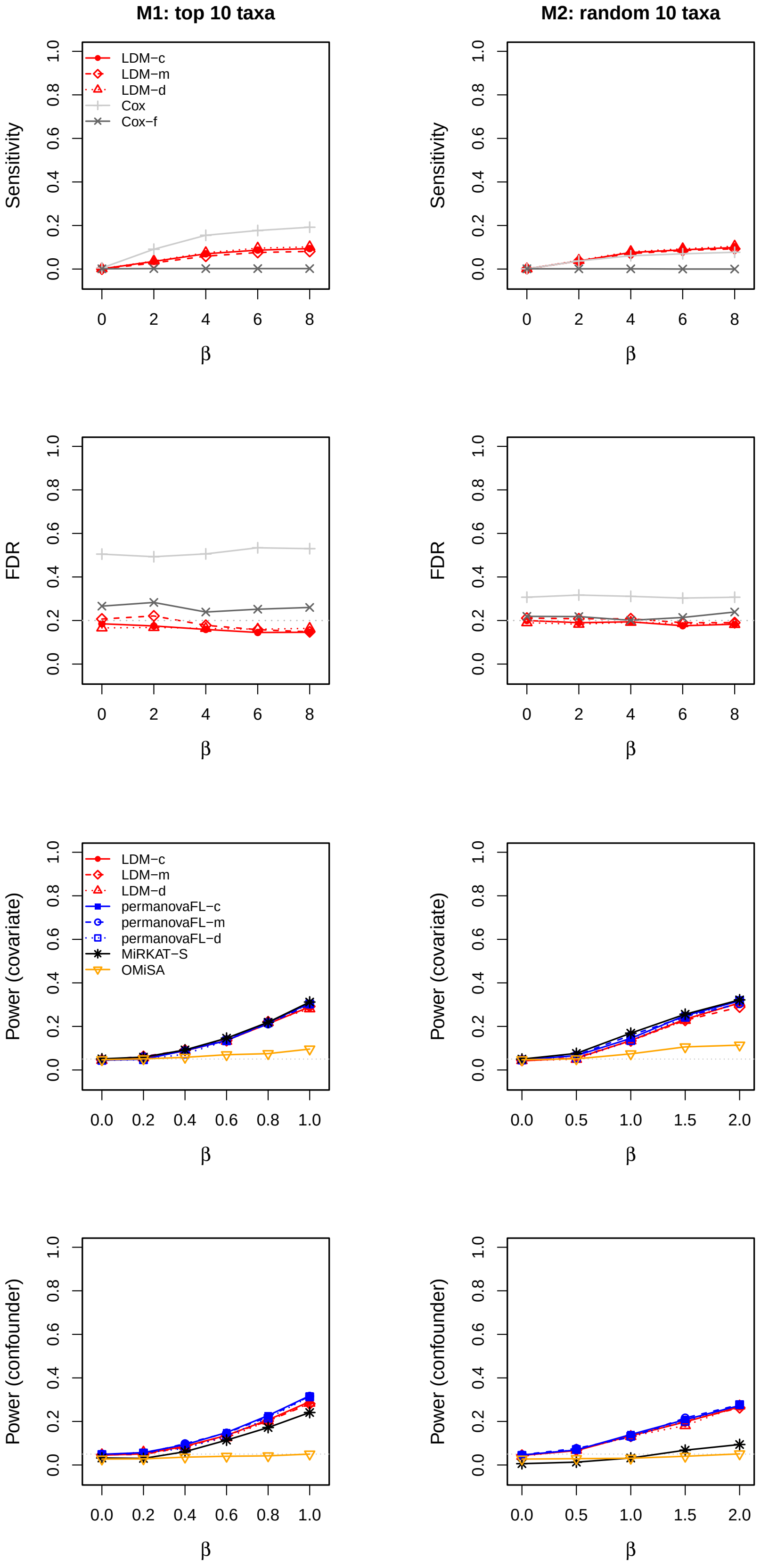


Figure S5: Results for simulated data with 50% censoring and $n = 50$. Results of sensitivity and empirical FDR were obtained when $X_i$ was a confounder ($\beta_{XZ} = 0.8$).

(a) Overall survival

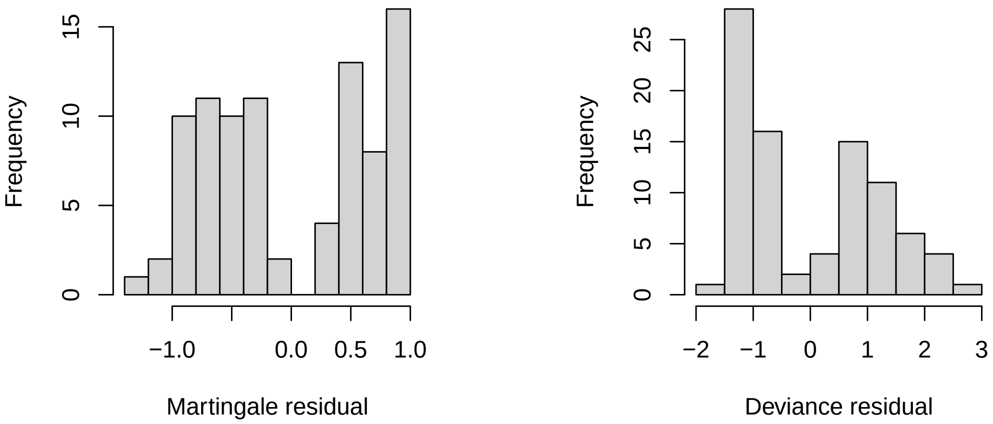


(b) Time to stage-III aGVHD

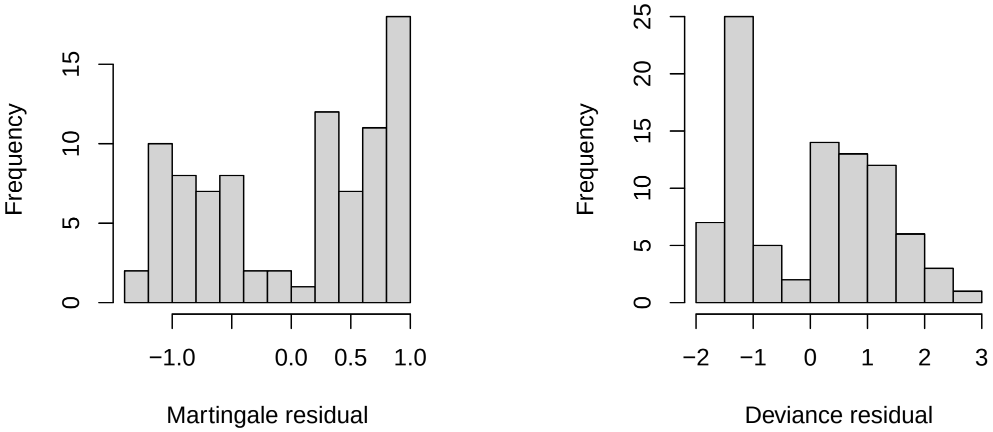


Figure S6: Martingale and deviance residuals, generated from the Cox model that fit age and gender as covariates in analysis of the aGVHD data.

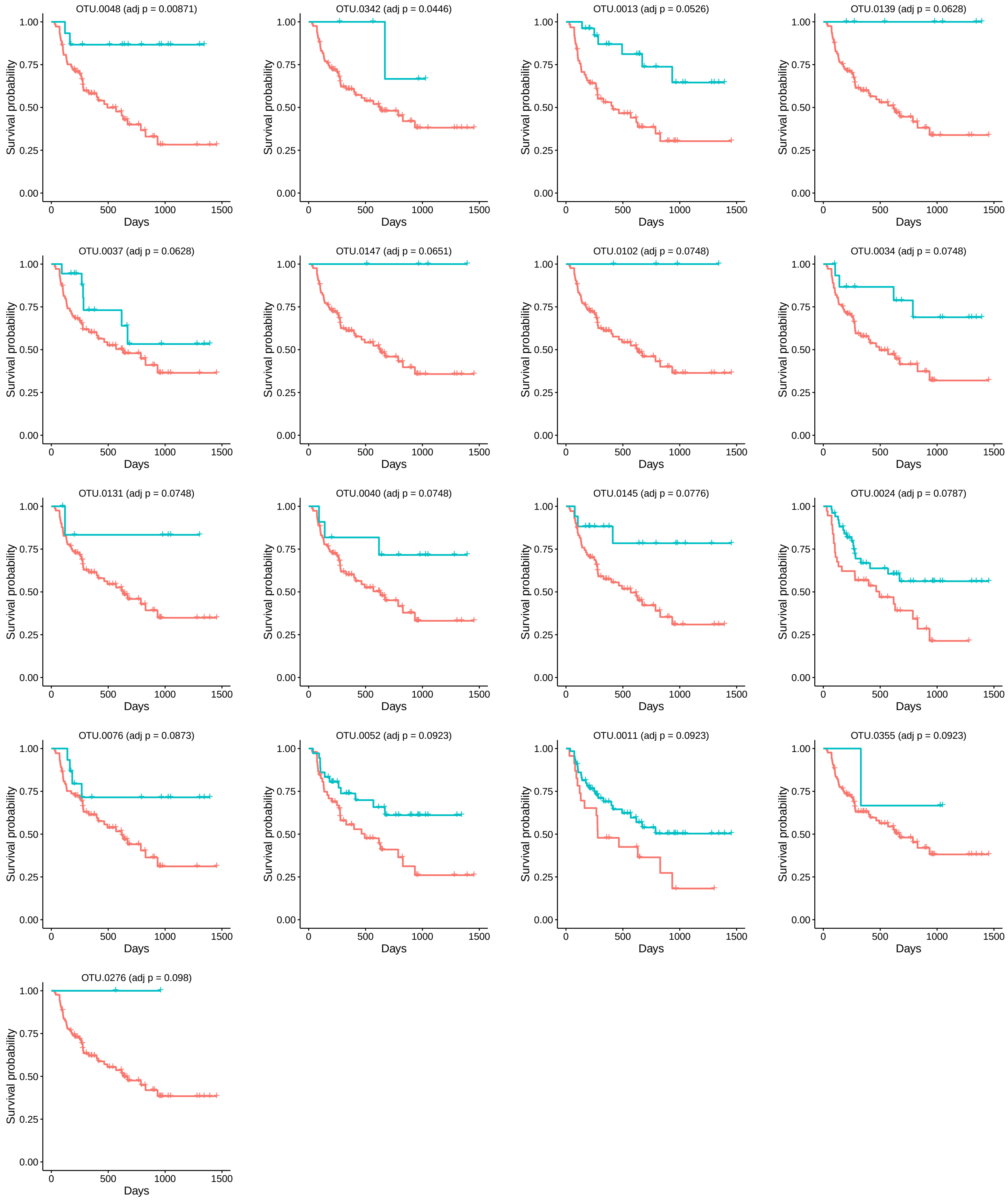


Figure S7: Survival functions for the overall survival outcome by the presence (blue) and absence (red) status (based on a singly rarefied OTU table) of the OTUs detected by LDM-c. The plots were ordered by the adjusted $p$-values from LDM-c.

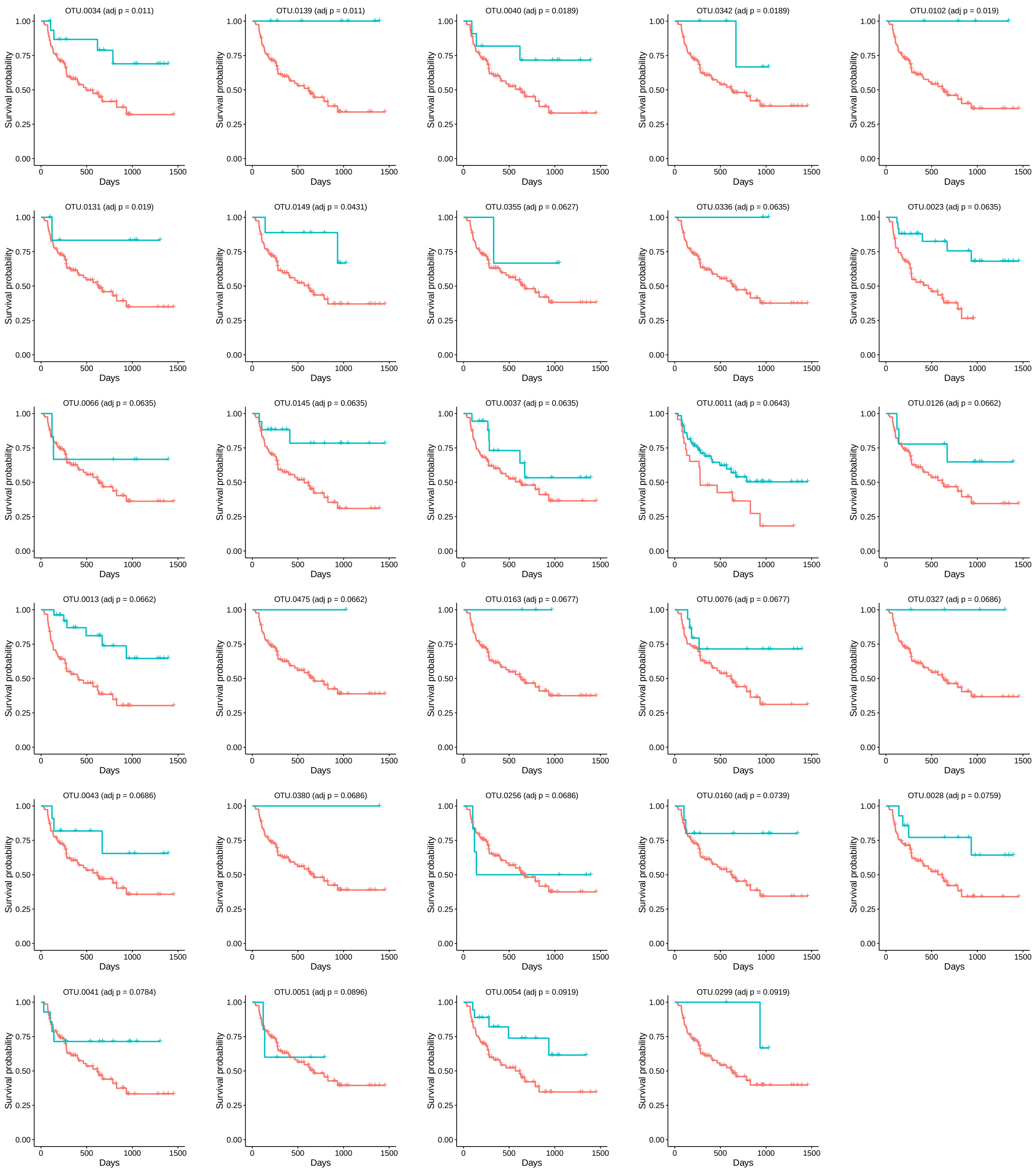


Figure S8: See the caption to Figure S7. The outcome is the time to stage-III aGVHD here.
